# IRP1 deficiency alters mitochondrial metabolism and protects against metabolic syndrome pathologies

**DOI:** 10.1101/2025.08.19.671129

**Authors:** Wen Gu, Nicole Wilkinson, Carine Fillebeen, Darren Blackburn, Korin Sahinyan, Eric Bonneil, Tao Zhao, Zhi Luo, Vahab Soleimani, Vincent Richard, Christoph H. Borchers, Albert Koulman, Benjamin Jenkins, Bernhard Michalke, Hans Zischka, Judith Sailer, Vivek Venkataramani, Othon Iliopoulos, Gary Sweeney, Kostas Pantopoulos

**Affiliations:** Lady Davis Institute for Medical Research, Jewish General Hospital and Department of Medicine, McGill University, Montreal, Quebec, Canada; Institute for Research in Immunology and Cancer, University of Montreal, Montreal, Quebec, Canada; Hubei Hongshan Laboratory, Fishery College, Huazhong Agricultural University, Wuhan 430070, China; Institute of Metabolic Science-Metabolic Research Laboratories, University of Cambridge, Cambridge, UK; Helmholtz Zentrum München GmbH – German Research Center for Environmental Health, Research Unit Analytical BioGeoChemistry, Neuherberg, Germany; Environmental Hygiene, Technical University Munich, Munich, Germany; Helmholtz Zentrum München GmbH – German Research Center for Environmental Health, Institute of Molecular Toxicology and Pharmacology, Neuherberg, Germany; Comprehensive Cancer Center Mainfranken, Julius-Maximilians Universität Würzburg, Germany; Department of Medicine, Hematology-Oncology Unit, Massachusetts General Hospital and the Massachusetts General Hospital Cancer Center, Boston, MA; Department of Biology, York University, Toronto, Ontario, Canada

**Keywords:** iron metabolism, mitochondrial dysfunction, gluconeogenesis, metabolic reprogramming, hepatic steatosis, MASLD

## Abstract

Iron regulatory protein 1 (IRP1) is a post-transcriptional regulator of cellular iron metabolism. In mice, loss of IRP1 causes polycythemia through translational de-repression of hypoxia-inducible factor 2α (HIF2α) mRNA, which increases renal erythropoietin production. Here we show that *Irp1^-/-^* mice develop fasting hypoglycemia and are protected against high-fat diet–induced hyperglycemia and hepatic steatosis. Discovery-based proteomics of *Irp1^-/-^* livers revealed a mitochondrial dysfunction signature. Seahorse flux analysis in primary hepatocytes and differentiated skeletal muscle myotubes confirmed impaired respiratory capacity, with a shift from oxidative phosphorylation to glycolytic ATP production. This metabolic rewiring was associated with enhanced insulin sensitivity and increased glucose uptake in skeletal muscle. Under metabolic stress, IRP1 deficiency altered the redox balance of mitochondrial iron, resulting in inefficient energy production and accumulation of amino acids and metabolites in skeletal muscle, rendering them unavailable for hepatic gluconeogenesis. These findings identify IRP1 as a critical regulator of systemic energy homeostasis.

## Introduction

Iron regulatory proteins, IRP1 and IRP2 coordinately control the expression of mRNAs containing iron responsive elements (IREs). These include among others *Tfrc*, *Fth/Ftl* and *Slc40α1* mRNAs, which encode proteins of iron uptake (transferrin receptor 1), storage (ferritin) and efflux (ferroportin), respectively (Galy *et al*, 2024). In iron-starved cells, IRE/IRP interactions protect *Tfrc* mRNA against degradation and inhibit translation of *Fth/Ftl* and *Slc40α1* mRNAs, in a homeostatic response to secure adequate iron supply. Conversely, in iron-replete cells, IRP1 is converted to cytosolic aconitase at the expense of its RNA-binding activity, while IRP2 undergoes proteasomal degradation. Inactivation of IRPs permits the decay of *Tfrc* mRNA and the *de novo* synthesis of ferritin and ferroportin to prevent detrimental iron overload.

IRP1 and IRP2 exhibit extensive sequence homology but only partial functional redundancy (Wilkinson & Pantopoulos, 2014). While global, or tissue-specific disruption of both IRPs has yielded lethal phenotypes (Galy *et al*, 2008; Galy *et al*, 2010; Smith *et al*, 2006), single *Irp1^-/-^* and *Irp2^-/-^* mice are viable and have distinct pathophysiological features. Thus, *Irp1^-/-^*mice develop polycythemia due to translational de-repression of the IRE-containing mRNA encoding HIF2α, which in turn stimulates erythropoiesis via transcriptional induction of erythropoietin in the kidney (Anderson *et al*, 2013; Ghosh *et al*, 2013; Wilkinson & Pantopoulos, 2013a). On the other hand, *Irp2^-/-^* mice manifest microcytic anemia, mild iron overload in the liver and duodenum, erythropoietic protoporphyria and late-onset neurodegeneration (Cooperman *et al*, 2005; Galy *et al*, 2005b).

Herein, we show that *Irp1^-/-^* mice are protected against hyperglycemia, insulin resistance and liver steatosis. These are common clinical manifestations of the metabolic syndrome, a pathologic state defined by the combined presentation of at least three of the following conditions: abdominal obesity, hyperglycemia due to insulin resistance, dyslipidemia and hypertension. Metabolic syndrome has a prevalence of ∼25% in the adult US population and increases the risk for type 2 diabetes, cardiovascular and liver disease, and all-cause mortality (Beltran-Sanchez *et al*, 2013). We demonstrate here that IRP1 deficiency causes mitochondrial dysfunction and metabolic rewiring that alters energy homeostasis. Our data uncover an unexpected role of IRP1 in the control of intermediary metabolism.

## Results

### Irp1^-/-^ mice are protected against high fat diet-induced hyperglycemia

Young *Irp1^-/-^* mice develop erythrocytosis due to induction of the HIF2α/erythropoietin axis (Anderson *et al*., 2013; Ghosh *et al*., 2013; Wilkinson & Pantopoulos, 2013a), and gradually recover after the age of 8-10 weeks, where HIF2α overexpression is mitigated (Wilkinson & Pantopoulos, 2013a). Prompted by evidence that HIF2α also controls insulin signaling in the liver via transcriptional induction of insulin receptor substrate 2 (IRS2) (Taniguchi *et al*, 2013; Wei *et al*, 2013), we analyzed metabolic functions in these animals. At the age of 5 weeks, *Irp1^-/-^*mice exhibited profound fasting hypoglycemia (Fig. 1A), while the phenotype of heterozygous *Irp1^+/-^* animals was indistinguishable from wild type controls (data not shown). Insulin tolerance testing (ITT) caused a sharper early drop in blood glucose levels of *Irp1^-/-^* mice (Fig. 1B), indicating increased insulin sensitivity. Moreover, *Irp1^-/-^* mice could not initiate a gradual return of blood glucose levels to baseline within 45-60 min post insulin injection; instead, they developed hypoglycemic seizures and had to be treated for recovery. In line with the severe fasting hypoglycemia, we observed 18% reduced survival of *Irp1^-/-^*pups at the age of 4 weeks (Fig. 1C), which was more pronounced in male (24%) vs female (11%) animals (Fig. S1A). Notably, blood glucose levels were normalized in older *Irp1^-/-^* mice (Fig. 1D, control diet), but the age-dependent correction of hypoglycemia was abolished when animals were fed an iron-deficient diet (IDD) (Fig. 1D). These data suggest that IRP1 deficiency has an impact on metabolism and are consistent with HIF2α involvement.

**Figure 1.**
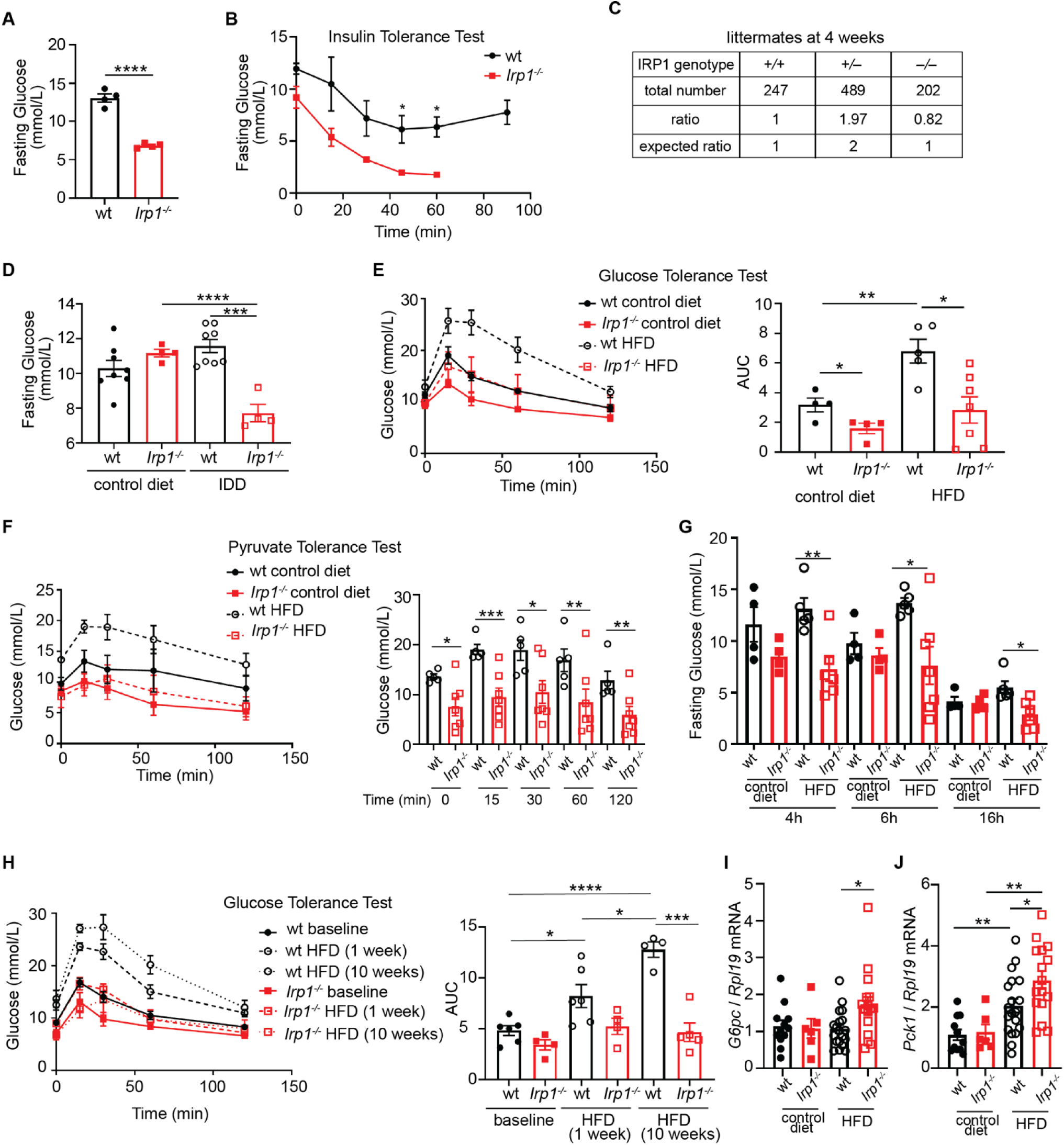
Young *Irp1^-/-^*mice are hypoglycemic and adult animals do not develop high fat diet-induced hyperglycemia. Male *Irp1^-/-^* and wild type mice (n=4-8 per experimental group) were analyzed at weaning (at the age of 5 weeks) or following dietary interventions initiated immediately after weaning (10-week long, unless otherwise indicated). (A) Fasting (5 h) blood glucose levels; and (B) insulin tolerance test (ITT) at weaning, after 4 h fasting. (C) Genotyping of all *Irp1^+/+^*, *Irp1^+/-^*and *Irp1^-/-^* littermates generated throughout this study. (D) Fasting (5 h) blood glucose levels in littermate mice fed a control or an iron-deficient diet (IDD). (E) Glucose tolerance test (GTT) after 5 h fasting; and (F) pyruvate tolerance test (PTT) after 6 h fasting in littermate mice fed a control or a high-fat diet (HFD). (G) Blood glucose levels in littermate mice fed control or HFD following fasting for 4, 6 or 16 h. (H) GTT in littermate mice after 4 h fasting at baseline, and after feeding HFD for 1 week and 10 weeks. (I and J) qPCR analysis of liver *G6pc* (I) and *Pck1* (J) mRNA expression in non-fasted littermate mice fed control or HFD for 12 weeks. The right panels in (E) and (H) show area under curve (AUC); and in (F) and (H) pairwise comparisons of data from HFD-fed wild type and *Irp1^-/-^* mice at various time points. Quantitative data are presented as the mean±SEM. Statistically significant differences are indicated by * (p<0.05), ** (p<0.01), *** (p<0.001), or **** (p<0.0001), respectively.

We then analyzed the responses of *Irp1^-/-^* mice and wild type littermates to a high-fat diet (HFD). Young (5-6 weeks old) male animals were fed a HFD or control diet for 10 weeks and, subsequently, were subjected to glucose tolerance testing (GTT). While, as expected, wild type mice on HFD developed hyperglycemia and glucose intolerance, *Irp1^-/-^* mice maintained physiological blood glucose levels and glucose clearance despite HFD feeding (Fig. 1E). The observed metabolic differences were independent of serum insulin levels (Fig. S1B) or weight gain (Fig. S1C), which were similar in HFD-fed wild type and mutant animals. No significant differences in blood pressure were noted among the experimental groups (Fig. S1D-E). We noted that the hypoglycemic phenotype persisted in both male and female young *Irp1^-/-^* mice (data not shown). However, female *Irp1^-/-^* mice appeared less protected than males against HFD-induced hyperglycemia (Fig. S1F). Thus, we focused on male animals for mechanistic studies.

Pyruvate tolerance testing (PTT) revealed that *Irp1^-/-^*mice exhibit a modest impairment in pyruvate-induced hepatic gluconeogenesis (Fig. 1F). This was corroborated by decreased glucose levels in these animals following 16 hours fasting (Fig. 1G), under conditions where glycogen stores are depleted and glucose supply largely depends on hepatic gluconeogenesis (Geisler *et al*, 2016). Furthermore, *Irp1^-/-^*mice on HFD exhibited lower blood glucose levels after 4 and 6 hours of fasting compared to wild type controls, indicating reduced hepatic glycogen storage (Fig. 1G). This hypoglycemic phenotype was also recapitulated in a longitudinal experiment, in which *Irp1^-/-^*mice and wild type littermates were subjected to GTT at baseline, and at 1 and 10 weeks after initiation of HFD intake. Unlike wild type animals, which developed increased glucose intolerance with prolonged HFD feeding, the *Irp1^-/-^* mice exhibited minimal defects in glucose clearance mostly after 1 week of HFD intake but not at the endpoint (10 weeks), suggesting metabolic adaptation and reprogramming (Fig. 1H). To gain mechanistic insights, we analyzed hepatic expression of genes involved in gluconeogenesis (*G6pc*, *Pck1*, *Fbp1*) in wild type and *Irp1^-/-^* mice fed control diet or HFD. *G6pc* (Fig. 1I) and *Pck1* (Fig. 1J) were significantly upregulated in *Irp1^-/-^* vs wild type animals on HFD, and a similar trend appeared for *Fbp1* (Fig. S1G), despite the overall impaired gluconeogenic capacity. The expression of HIF2α-inducible *Irs2* (but not *Irs1*) tended to be higher in *Irp1^-/-^*mice on HFD (Fig. S1H). This indicates potentially increased insulin sensitivity and is again consistent with a role of HIF2α. The induction of gluconeogenic genes despite increased insulin sensitivity suggests that *Irp1^-/-^*mice are in an energy deprived state.

Metabolic cage analysis of wild type and *Irp1^-/-^* mice on HFD showed no significant differences among the genotypes on food or water intake, oxygen consumption, CO_2_ production, respiratory exchange rate or heat, while *Irp1^-/-^* mice tended to exhibit slightly increased movement activities (Fig. S2).

### Irp1^-/-^ mice are protected against high fat diet-induced liver steatosis

After 10 weeks of HFD feeding, livers from wild type mice became pale, indicative of steatosis; by contrast, livers from *Irp1^-/-^*littermates retained their physiological color, suggesting protection against steatosis (Fig. 2A). Histopathological analysis using hematoxylin and eosin (H&E) or oil red O (ORO) staining confirmed that liver sections from *Irp1^-/-^* mice had significantly reduced fat content, without large visible lipid droplets (Fig. 2B), indicating that these animals were spared from HFD-induced hepatic steatosis.

**Figure 2.**
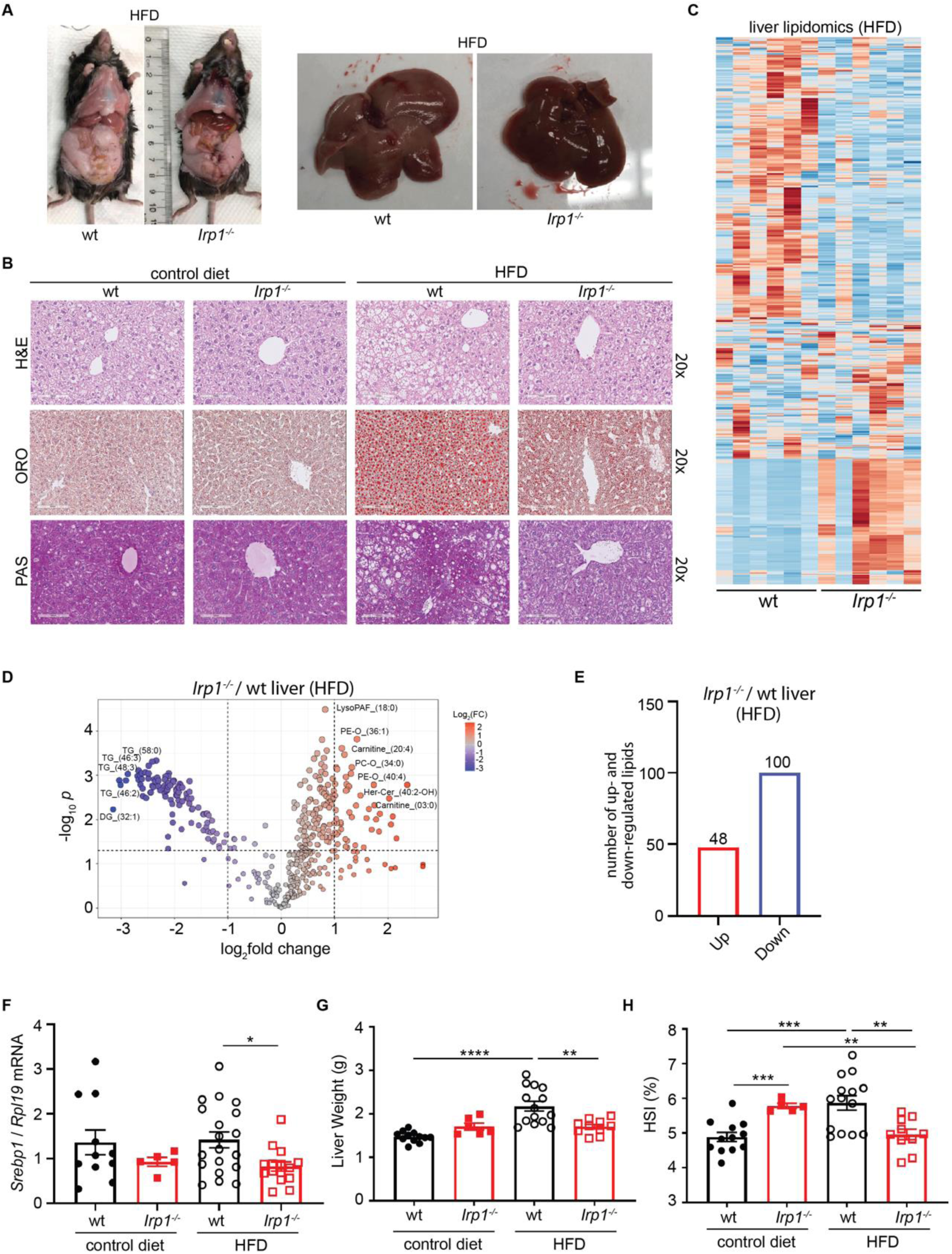
*Irp1^-/-^*mice are protected against high fat diet-induced liver steatosis. 5-weeks old male *Irp1^-/-^* mice and wild type littermates (n=6-12 per experimental group) were provided a control or a high-fat diet (HFD) for 10 weeks. At the endpoint, the mice were euthanized, and livers were dissected for histological and biochemical analysis. (A) Images of representative mice and dissected livers after euthanasia. (B) Histopathological analysis of liver sections by staining with H&E for tissue architecture (top), oil red O (ORO) for neutral fats and lipids (middle) and Periodic Acid-Schiff (PAS) for glycogen (bottom); original magnification: 20x. (C) Heatmap of liver lipidomic changes between wild type and *Irp1^-/-^* mice on HFD (n=6 mice from each genotype). (D) Volcano plot of differentially expressed lipids in the liver of *Irp1^-/-^* and wild type mice on HFD; p value threshold was set to 0.05 and fold change to 2. (E) Number of up- and down-regulated lipids in the liver of *Irp1^-/-^* vs wild type mice on HFD. (F) qPCR analysis of liver *Srebp1* mRNA expression. (G) Liver weight. (H) Hepatosomatic index (HSI). Quantitative data are presented as the mean±SEM. Statistically significant differences are indicated by ** (p<0.01), *** (p<0.001), or **** (p<0.0001), respectively.

The histological findings were corroborated by large-scale lipidomics profiling; the heatmap in Fig. 2C reveals dramatic alterations in the liver lipidome of *Irp1^-/-^* mice. A total of 100 lipid species were upregulated and 48 downregulated in the liver of *Irp1^-/-^* vs wild type mice (Fig. 2D-E). Upregulated lipids include phosphatidylethanolamine plasmalogens (PE-O_(36:1) and PE-O_(40:4)), arachidonoyl-L-carnitine (carnitine_(20:4)), and phosphatidylcholine plasmalogens (PC-O_(34:0) and PCO_(31:0)). Downregulated lipids include triacylglycerols (TG_(58:0), TG_(46:3), TG_(48:3) and TG_(46:2), and diacylglycerols (DG_(32:1), which may indicate decreased hepatic lipogenesis in *Irp1^-/-^* mice. Supporting evidence is provided by the significantly reduced expression of *Srebp1* mRNA (Fig. 2F) encoding a key transcriptional activator of *de novo* hepatic lipogenesis (Wakil & Abu-Elheiga, 2009).

Consistent with these findings, HFD feeding increased liver weight and the hepatosomatic index (liver weight expressed as a percentage of total body weight) in wild type mice only (Fig. 2G-H). In contrast, the hepatosomatic index decreased in HFD-fed *Irp1^-/-^* animals. Periodic Acid-Schiff (PAS) staining suggested that livers from *Irp1^-/-^* mice on HFD also had reduced glycogen content (Fig. 2B, bottom panel), in line with the relative hypoglycemia that these animals develop after 4 and 6 hours fasting (Fig. 1G).

Serum biochemistry showed that HFD increased total cholesterol in both wild type and mutant mice (Fig. S3A), while triglyceride levels tended to be lower in *Irp1^-/-^*mice independently of diet (Fig. S3B). Despite elevated HDL cholesterol in both genotypes following HFD intake (Fig. S3C), non-HDL cholesterol levels appeared lower in *Irp1^-/-^* animals (Fig. S3D) and total cholesterol/HDL ratios were not affected (Fig. S3E). Values for serum iron, total iron-binding capacity (TIBC), transferrin saturation and liver *Hamp* mRNA did not differ among all experimental groups (Fig. S3F-I), while serum ferritin was elevated in *Irp1^-/-^* mice (Fig. S3J), possibly due to translational de-repression under IRP1 deficiency. Complete blood count showed a normal red blood cell number and even reduced hematocrit in *Irp1^-/-^*mice after the 10-week HFD feeding period (Table S1), validating the correction of polycythemia with age (Wilkinson & Pantopoulos, 2013b). Along these lines, liver *Epo* mRNA and serum Epo levels were similar in all groups (Fig. S3K-L). Nevertheless, splenomegaly persisted in the *Irp1^-/-^*animals (Fig. S3M-O), indicating extramedullary hematopoiesis. This was not associated with altered splenic iron content (Fig. S3P).

### IRP1 deficiency causes mitochondrial dysfunction and a shift of energy metabolism from oxidative phosphorylation to glycolysis

An unbiased proteomics approach was utilized to elucidate pathways by which IRP1 deficiency affects metabolic functions. To this end, liver samples from wild type and *Irp1^-/-^* mice on HFD (n=4 from each group) were subjected to LC-MS/MS analysis. After filtering, on average 2300 proteins were identified in each sample. Principal component analysis shows good separation between samples from the two genotypes (Fig. S4A), while Pearson correlation analysis shows that the replicates correlate well (Fig. S4B). 101 proteins were up-regulated and 82 down-regulated in *Irp1^-/-^* vs wild type livers (Fig. 3A-B). Up-regulated proteins include S100a1, Slc4a1, Blvra etc. Prominent down-regulated proteins were Nduf3, Adcy5, Prdm16 and IRP1 (Aco1), which serves as control. Enrichment analysis with IPA software identified mitochondrial pathways (oxidative phosphorylation, electron transport, ATP synthesis) as mostly affected by IRP1 deficiency in the liver of HFD-fed mice (Fig. 3C). The top hits in GO hierarchy analysis were catalytic activity (GO Molecular Function), small molecule metabolic process (GO Biological Process), cytoplasm and mitochondrion (GO Cellular Component), while the top hits in KEGG and Reactome pathway analysis were metabolic pathways and metabolism, respectively (Fig. S5). These data uncover metabolic reprograming in the liver of *Irp1^-/-^* mice that involves mitochondria. Mitochondrial dysfunction is also highlighted in STRING functional association networks (Fig. 3D) and validated by Western blot analysis for electron transport chain complexes, showing reduced expression especially of proteins representing complexes I and II, but also III and IV in the liver of *Irp1^-/-^* mice (Fig. 3E). This was not accompanied by any significant changes in expression of markers of mitochondrial biogenesis (*Pgc1α* mRNA), fission (*Fis1* and *Drp1* mRNAs) or fusion (*Mfn1* and *Mfn2* mRNAs) (Zhao *et al*, 2024) (Fig. S6A-E).

**Figure 3.**
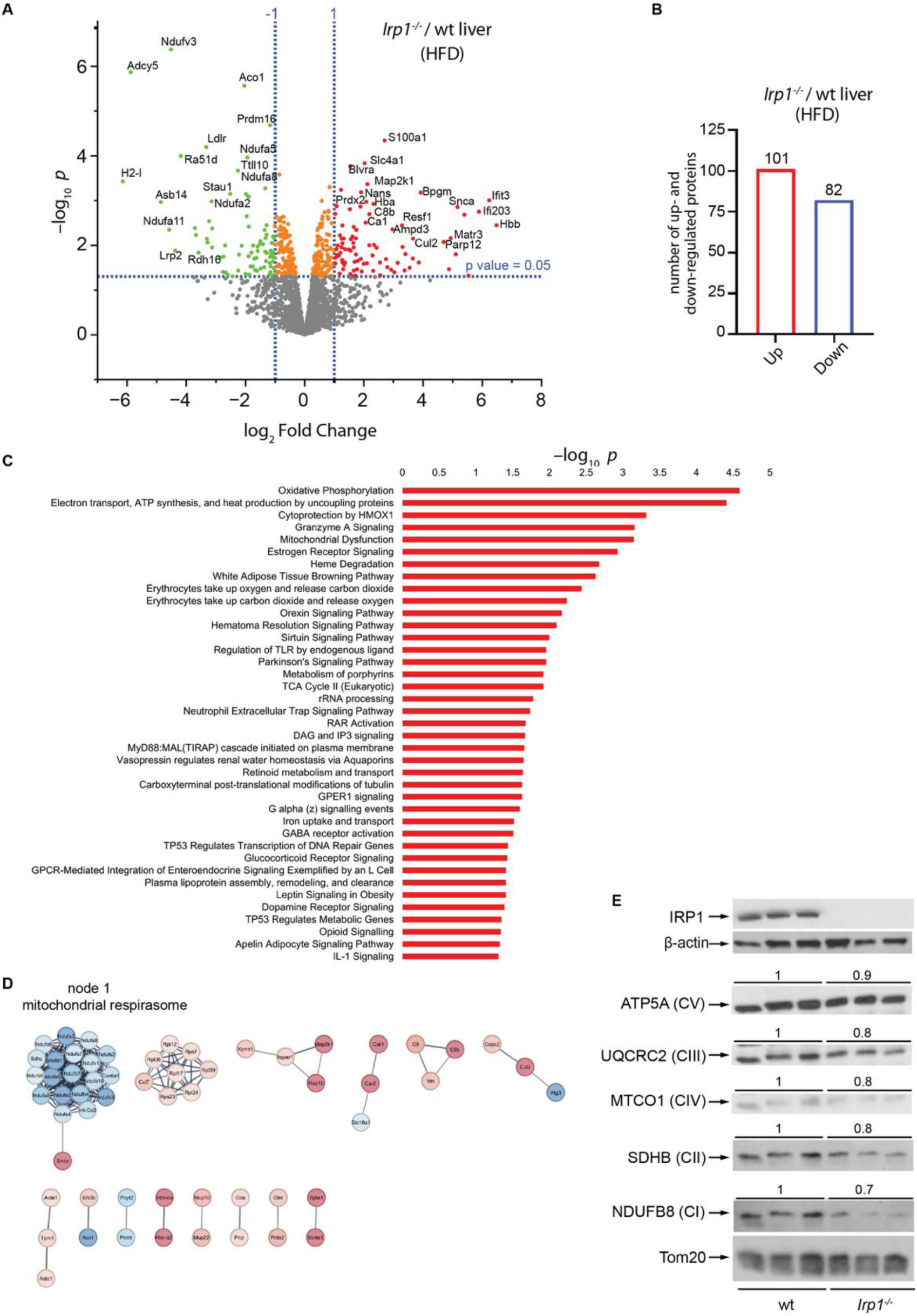
Altered mitochondrial proteome in the liver of *Irp1^-/-^* mice. 5-weeks old male *Irp1^-/-^* mice and wild type littermates (n=4 per experimental group) were provided a high-fat diet (HFD) for 10 weeks. At the endpoint, the mice were euthanized, and livers were dissected for proteomics analysis. (A) Volcano plot of differentially expressed proteins (*Irp1^-/-^* vs wild type); p value threshold was set to 0.05 and fold change to 2. (B) Number of up- and down-regulated proteins in the liver of *Irp1^-/-^* vs wild type mice. (C) Pathway enrichment analysis generated by the Qiagen IPA software. (D) STRING networks based on all differentially expressed proteins show that the major enrichment in GO terms and KEGG pathways is linked to mitochondrial respirasome. (E) Western blot analysis of mitochondrial respiratory chain proteins corresponding to complexes I, II, III, IV and V using whole liver extracts from three representative mice in each group. The membranes were also probed with antibodies against IRP1, β-actin (cytosolic loading control) and Tom20 (mitochondrial loading control). Blots were quantified by densitometry relative to Tom20 and average values are shown on top. Ratios of respiratory chain proteins to Tom20 from wild type mice are set as 1.

Diminished expression of electron transport chain complexes has previously been reported in *Irp1^-/-^* murine embryonic fibroblasts (MEFs) (Li *et al*, 2018). To assess functional implications, we analyzed mitochondrial oxygen consumption rates in *Irp1^-/-^* MEFs with the Seahorse assay. IRP1 deficiency significantly impaired basal, ATP-linked and maximal respiration, as well as spare capacity (Figs. 4A and S7A-D). This was associated with increased glycolytic activity and lactate production (Fig. 4B-C); both basal and compensatory glycolysis were induced (Fig. S7E-F). Similar results were obtained with differentiated primary skeletal muscle myotubes from *Irp1^-/-^* mice (Figs. 4D-E and S7G-L), which also exhibited defective mitochondrial respiration when palmitate was used as substrate (Figs. 4F and S7M-P). Along these lines, muscle fibers from *Irp1^-/-^* mice had increased glucose uptake capacity (Fig. 4G). IRP1 deficiency did not affect muscle stem cell differentiation to myotubes (Fig. S8A) but increased proliferation rates of myoblasts (Fig. S8B) and MEFs (Fig. S8C). In line with the liver proteomics data, primary hepatocytes from *Irp1^-/-^* mice likewise exhibited impaired mitochondrial respiration (Fig. 4H and S7Q-T) and increased basal glycolysis (Fig. 4I). The effects in hepatocytes appeared milder compared to MEFs and myotubes, and statistically significant differences among the genotypes were only noted for spare capacity (Fig. S7T) and basal (Fig. 4I), but not compensatory glycolysis (not shown). Taken together, the above data suggest that IRP1 deficiency triggers mitochondrial dysfunction and a shift to aerobic glycolysis for energy production.

**Figure 4.**
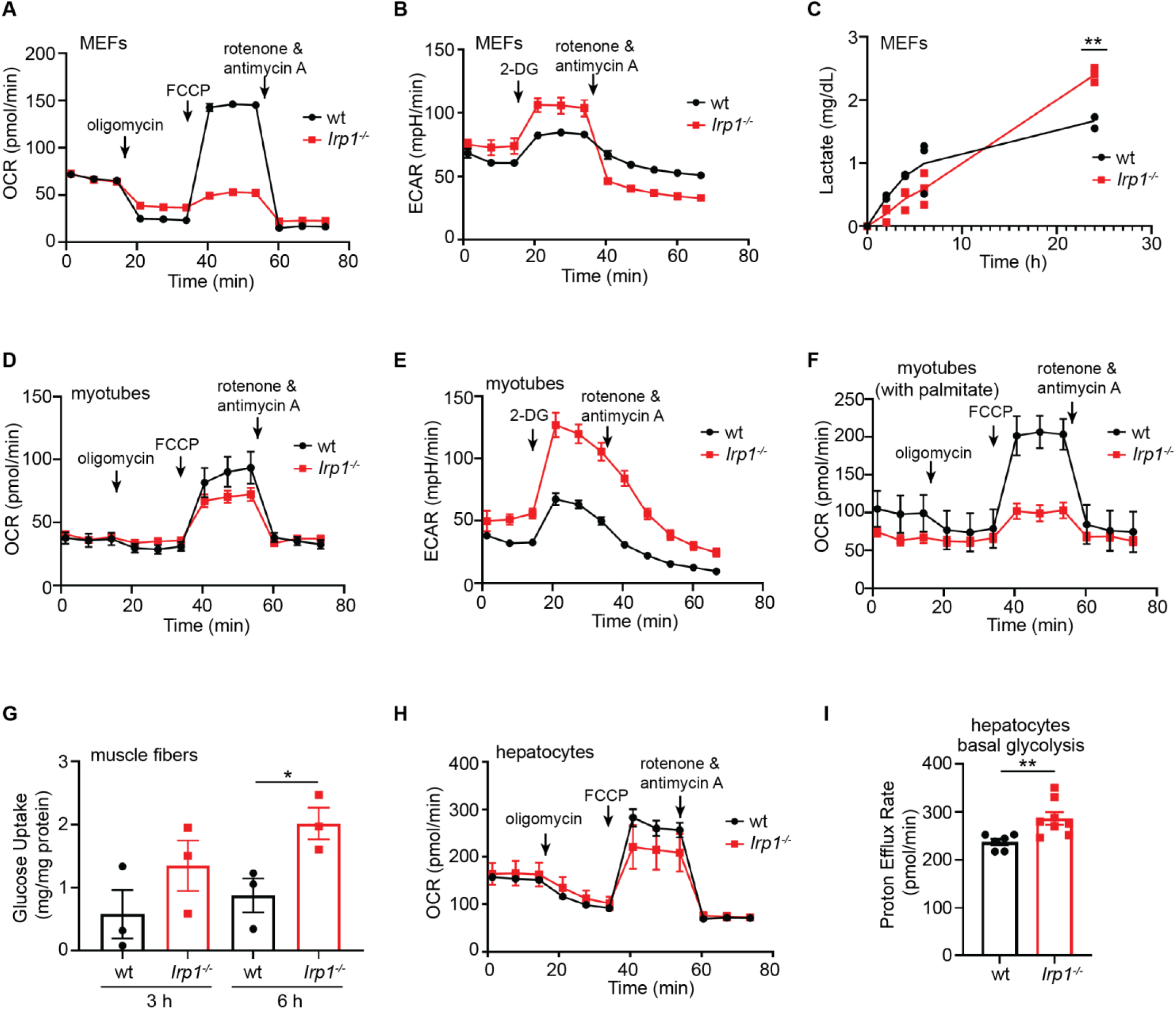
IRP1 deficiency causes mitochondrial dysfunction and a switch from oxidative energy metabolism to aerobic glycolysis. Mouse embryonic fibroblasts (MEFs), primary hepatocytes and differentiated myotubes from *Irp1^-/-^* and wild type mice were analyzed for mitochondrial function by the Seahorse assay. The time for addition of oligomycin, FCCP and rotenone/antimycin A or 2-deoxy-glucose (2-DG) is indicated by arrows. (A) Oxygen consumption rates (OCR) in *Irp1^-/-^* and wild type MEFs. (B) Extracellular acidification rates (ECAR) in *Irp1^-/-^*and wild type MEFs. (C) Lactate production by *Irp1^-/-^* and wild type MEFs. (D) OCR in *Irp1^-/-^* and wild type differentiated myotubes. (E) ECAR in *Irp1^-/-^* and wild type differentiated myotubes. (F) OCR with palmitate as substrate in *Irp1^-/-^* and wild type differentiated myotubes. (G) Glucose uptake assay in muscle fibers from *Irp1^-/-^* and wild type mice. (H) OCR in *Irp1^-/-^* and wild type primary hepatocytes. (I) Proton efflux rate indicating basal glycolysis in *Irp1^-/-^* and wild type primary hepatocytes. Data in (G) and (I) are presented as the mean ± SEM; statistically significant differences are indicated by * (p<0.05) or ** (p<0.01).

### IRP1 deficiency alters mitochondrial redox speciation of iron in the liver and skeletal muscles

In mouse (Padda *et al*, 2015) and rat (Dongiovanni *et al*, 2015) models of liver steatosis, HFD intake elicited a phenotype of low hepatic iron characterized by induction of transferrin receptor 1 (TfR1), the major iron uptake protein. In agreement with the earlier findings, we noted that wild type mice on HFD manifested increased TfR1 levels in the liver (Fig. 5A). TfR1 expression was reduced, and ferritin was upregulated in the liver of *Irp1^-/-^*vs wild type mice on control diet, presumably due to IRP1 deficiency. These responses were preserved following HFD feeding; however, a small number (<25%) of the analyzed livers from *Irp1^-/-^*animals exhibited high TfR1 and low ferritin levels. On the other hand, HFD intake neither induced TfR1 nor suppressed ferritin in skeletal muscles of wild type mice (Fig. 5B), indicating relatively normal iron balance in this tissue. Ferritin was paradoxically suppressed in skeletal muscles of HFD-fed *Irp1^-/-^*mice and TfR1 levels appeared slightly reduced, possibly due to IRP1 deficiency. These data suggest that iron homeostasis in the liver and skeletal muscles are controlled by distinct tissue-specific pathways in response to HFD, with or without IRP1 deficiency.

**Figure 5.**
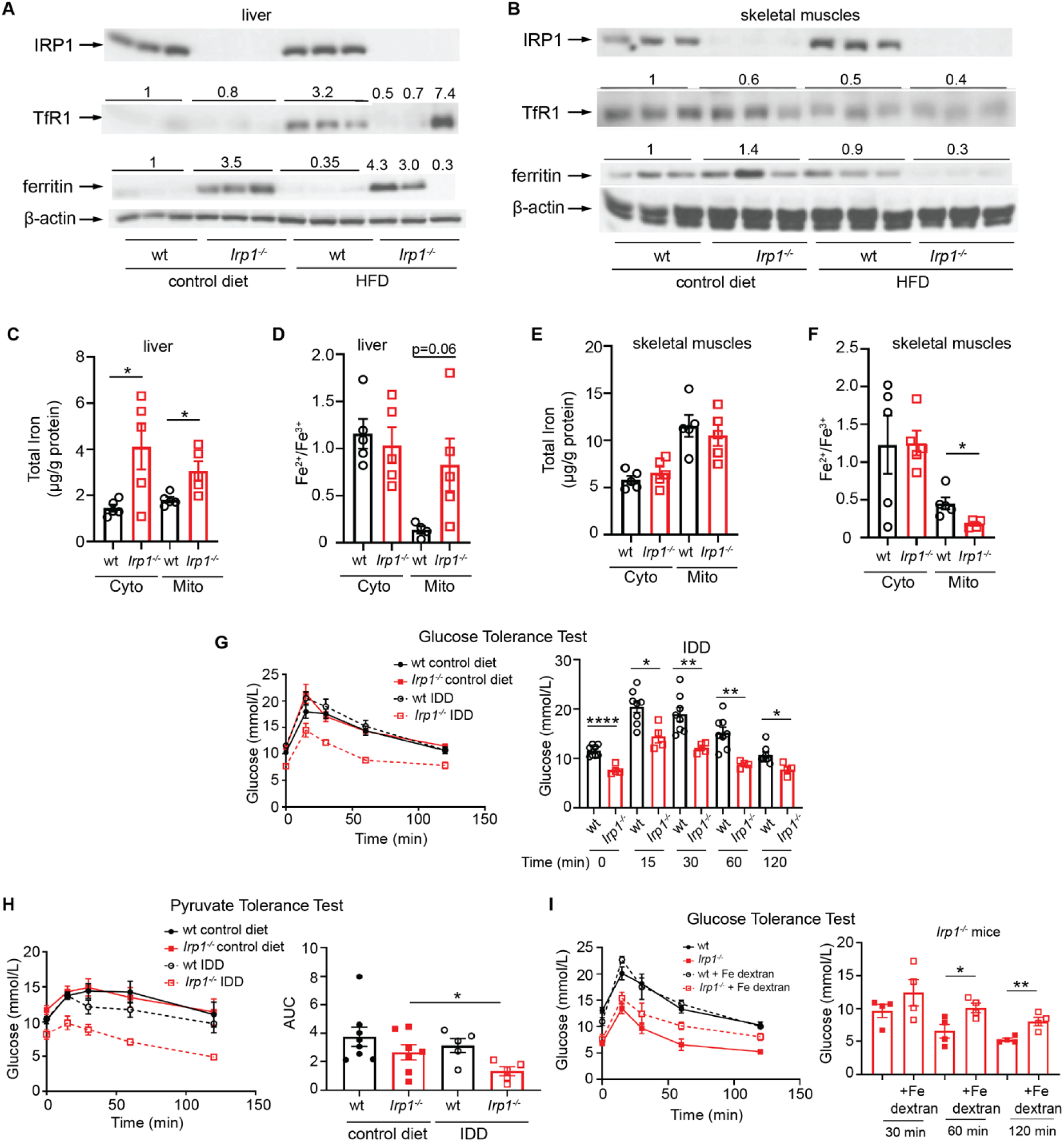
Evidence that mitochondrial disfunction in IRP1-deficient cells is caused by impaired mitochondrial redox speciation of iron. (A-F) 5-weeks old male *Irp1^-/-^* mice and wild type littermates (n=5 per experimental group) were provided a control or a high-fat diet (HFD) for 10 weeks. At the endpoint, the mice were euthanized, and liver and quadriceps skeletal muscle tissue samples were isolated for biochemical studies. Cytosolic and mitochondrial fractions were prepared and used for quantification of total iron and iron speciation analysis. (A-B) Western blot analysis of IRP1, TfR1, ferritin and β-actin in the liver (A) and skeletal muscles (B). (C-D) Quantification of total iron levels (C) and Fe^2+^/Fe^3+^ ratios (D) in cytosolic and mitochondrial fractions of the liver. (E-F) Quantification of total iron levels (E) and Fe^2+^/Fe^3+^ ratios (F) in cytosolic and mitochondrial fractions of skeletal muscles. (G-I) 5-weeks old male *Irp1^-/-^* mice and wild type littermates (n=5-7 per experimental group) were provided a control or an iron-deficient diet (IDD) for 10 weeks. (G) Glucose tolerance test (GTT) after 5 h fasting; the right panel shows pairwise comparisons of data from wild type and *Irp1^-/-^*mice on IDD at various time points. (H) Pyruvate tolerance test (PTT) after 6 h fasting; the right panel shows AUC. (I) 5-weeks old male *Irp1^-/-^* mice were subjected to GTT following injections with iron dextran or not, and after 5 h fasting; the right panel shows pairwise comparisons of data from *Irp1^-/-^* mice with or without iron dextran injections at various time points. In (A-B), Western blots for TfR1 and ferritin were quantified by densitometry relative to β-actin; average or individual (due to experimental variability) values are shown on top. TfR1/ β-actin and ferritin/ β-actin ratios from wild type mice are set as 1. In (C-F), iron measurements were normalized to protein content and are expressed as mean ± SD. Quantitative data in (C-I) are presented as the mean±SEM. Statistically significant differences are indicated by * (p<0.05), ** (p<0.01), or **** (p<0.0001), respectively.

To shed light into these pathways and dissect the role of IRP1, we profiled ferrous (Fe²⁺) and ferric (Fe³⁺) iron content in the cytosolic and mitochondrial fractions from liver and skeletal muscles of wild type and *Irp1^-/-^*mice on HFD, using cation exchange chromatography coupled with inductively coupled plasma mass spectrometry (SCX-ICP-MS). The Fe²⁺/Fe³⁺ ratio was calculated as an indicator of redox-active, labile iron pools. Purity of subcellular fractions was assessed by Western blotting for cytosolic IRP1 and mitochondrial Tom20 (Fig. S9A-B).

Total (Fe²⁺ and Fe³⁺) iron levels were significantly higher in the liver cytosolic (∼2.8-fold) and mitochondrial (∼1.7-fold) fractions of *Irp1^-/-^* mice compared to wild type controls (Fig. 5C). Separate measurements of Fe²⁺ and Fe³⁺ are shown in Fig. S9C-D. While cytosolic Fe²⁺/Fe³⁺ ratios were similar between genotypes, the mitochondrial Fe²⁺/Fe³⁺ ratio was elevated by ∼6-fold in *Irp1^-/-^* livers (Fig. 5D), suggesting a relative accumulation of reduced, potentially unutilized and redox-active Fe²⁺ in IRP1-deficient liver mitochondria.

In skeletal muscles, total iron (Fig. 5E), Fe²⁺ (Fig. S9E), Fe³⁺ (Fig. S9F) levels, and cytosolic Fe²⁺/Fe³⁺ ratios (Fig. 5D, right) were comparable across genotypes. However, mitochondrial Fe²⁺/Fe³⁺ ratios were significantly decreased ∼2.5-fold in *Irp1^-/-^* skeletal muscles (Fig. 5F), indicating a relative oxidation of the mitochondrial iron pool in the absence of IRP1. Collectively, these iron speciation data suggest that IRP1 deficiency impairs mitochondrial iron homeostasis in both liver and skeletal muscles under metabolic stress induced by HFD, while cytosolic iron buffering remains unaffected.

Contrary to data previously reported in MEFs (Li *et al*., 2018), IRP1 deficiency (or HFD intake) did not significantly affect expression of the Fe-S cluster biogenesis proteins frataxin and IscU in the liver and skeletal muscles (Fig. S9G-H).

Our findings suggest that the metabolic phenotype of *Irp1^-/-^*mice on HFD may be linked to defective mitochondrial iron homeostasis. Based on this, we hypothesized that dietary iron restriction might elicit metabolic responses in these animals analogous to those induced by HFD. Indeed, adult *Irp1^-/-^* mice fed an iron-deficient diet (IDD) not only maintained fasting hypoglycemia (Fig. 1D), but also exhibited improved glucose clearance (Fig. 5G) and blunted gluconeogenesis (Fig. 5H). Notably, injection with iron dextran did not affect glucose clearance in young (5-weeks old) wild type mice but substantially impaired it in hypoglycemic *Irp1^-/-^*littermates (Fig. 5I), underlying the critical role of iron in glucose metabolism. These data demonstrate that dietary iron deficiency mimics effects of metabolic stress imposed by HFD on glucose homeostasis in the absence of IRP1. Moreover, they support the notion that disruption of mitochondrial iron balance may be a common underlying factor contributing to fasting hypoglycemia in young *Irp1^-/-^* mice and to the metabolic responses of adult *Irp1^-/-^* mice to both HFD and IDD.

### Metabolic reprogramming in Irp1^-/-^ mice

Targeted metabolomics analysis in serum, liver and skeletal muscles was performed to uncover systemic metabolic alterations triggered by IRP1 deficiency following HFD or IDD intake. A total of 78 metabolites was detected in the serum, liver, and skeletal muscles of wild type and *Irp1^-/-^* mice on HFD; a complete heatmap is shown in Fig. S10A. Under these conditions, *Irp1^-/-^* mice exhibited lower levels of most serum and liver metabolites compared to wild type controls (Fig. 6A). We focused on changes in expression of specific metabolites such as glucogenic amino acids, ketogenic amino acids, gluconeogenesis precursors, fatty acids and TCA cycle intermediates (Fig. 6B). Among these, only aspartic acid was upregulated in serum, and citric and isocitric acid in the liver of *Irp1^-/-^* mice. By contrast, skeletal muscles from *Irp1^-/-^* mice on HFD were enriched in glucogenic amino acids and TCA cycle intermediates, with highest abundance of glutamine, proline and 2-ketoglutarate; expression of ketogenic lysine was also high. Enrichment analysis identifed the pathways most significantly affected by IRP1 deficiency and HFD. These include: a) Tryptophan metabolism and starch and sucrose metabolism in serum (Fig. S10B); glyoxylate and dicarboxylate metabolism, alanine aspartate and glutamate metabolism, tricarboxylic acid (TCA) cycle and unsaturated fatty acid biosynthesis in the liver (Fig. S10C); and c) arginine and proline metabolism, pantothenate and CoA biosynthesis, and pyrimidine metabolism in skeletal muscles (Fig. S10D).

**Figure 6.**
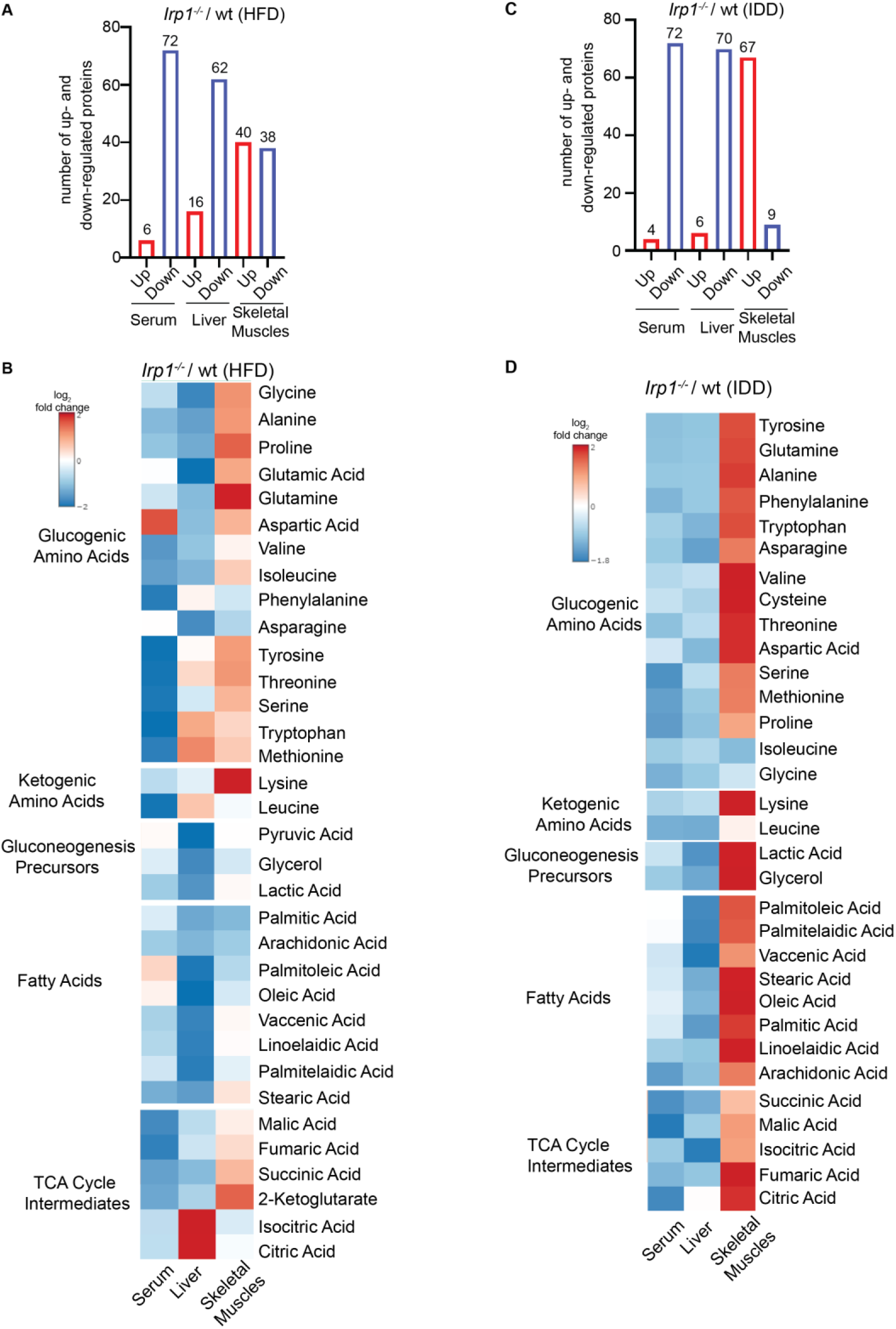
IRP1 deficiency triggers metabolic reprogramming in mice fed high-fat or iron-deficient diets. 5-weeks old male *Irp1^-/-^* mice and wild type littermates (n=4-5 per experimental group) were provided a high-fat diet (HFD) or an iron-deficient diet (IDD) for 10 weeks. At the endpoint, the mice were euthanized, and serum, liver and skeletal muscle samples were collected and processed for targeted metabolomics analysis. (A-B) Number of up- and down-regulated metabolites in serum, liver, and skeletal muscle of *Irp1^-/-^* vs wild type on HFD (A); and heatmap of detected glucogenic amino acids, ketogenic amino acids, gluconeogenesis precursors, fatty acids and TCA cycle intermediates (B). (C-D) Number of up- and down-regulated metabolites in serum, liver, and skeletal muscle of *Irp1^-/-^* vs wild type on IDD (C); and heatmap of detected glucogenic amino acids, ketogenic amino acids, gluconeogenesis precursors, fatty acids and TCA cycle intermediates (D). In (B) and (D), log_2_ fold changes are indicated on the left.

Under conditions of dietary iron restriction, the abundance of most analyzed metabolites was likewise lower in the serum and liver of *Irp1^-/-^* vs wild type mice (Fig. 6C-D and S11A). Notably, IDD intake resulted in a dramatic shift of metabolites to skeletal muscles of these animals. Thus, 67 out of 76 detected metabolites were upregulated in *Irp1^-/-^* vs wild type skeletal muscles, including glucogenic amino acids, gluconeogenesis precursors, fatty acids and TCA cycle intermediates. Pathways most significantly affected by IRP1 and iron deficiency include: a) Lipoic acid metabolism, butanoate metabolism, as well as alanine, aspartate and glutamate metabolism in serum (Fig. S11B); starch and sucrose metabolism, and tryptophan metabolism in the liver (Fig. S11C); and TCA cycle, glyoxylate and dicarboxylate metabolism, and biotin metabolism in skeletal muscles (Fig. S11D).

We also analyzed the impact of IRP1 deficiency under physiological conditions, in mice fed control diet. In general, metabolite abundance was lower in *Irp1^-/-^* vs wild type serum, liver or skeletal muscles (Fig. S12A-B). Enrichment analysis identified following mostly affected pathways: a) Glyoxylate and dicarboxylate metabolism, alanine, aspartate and glutamate metabolism, and glycolysis/gluconeogenesis in serum (Fig. S12C); histidine, metabolism, propanoate metabolism, pyruvate metabolism, TCA cycle, glyoxylate and dicarboxylate metabolism in the liver (Fig. S12D); and glycerophospholipid, fructose and mannose metabolism, inositol metabolism, glycerolipid metabolism and glycolysis/gluconeogenesis in skeletal muscles (Fig. S12E). The above data reveal extensive metabolic reprogramming in response to IRP1 deficiency, which is reflected in serum, liver, skeletal muscle metabolites. HFD or IDD intake trigger a shift of metabolites from liver and serum to skeletal muscles of *Irp1^-/-^*mice.

### IRP1 deficiency improves insulin sensitivity in hepatocytes and skeletal muscle cells

We explored whether the observed metabolic reprogramming in *Irp1^-/-^*mice is associated with altered insulin signaling. To this end, primary hepatocytes from wild type and *Irp1^-/-^* mice remained untreated or treated with insulin. Insuling signaling was monitored by analyzing Akt phosphorylation as a surrogate marker (Fig. 7A, left). p-Akt/Akt ratios were highly increased in insulin-treated *Irp1^-/-^*hepatocytes compared to insuling-treated wild type hepatocytes (Fig. 7A, right). Moreover, p-Akt/Akt ratios remained higher in *Irp1^-/-^* vs wild type hepatocytes following treatment with palmitate, a fatty acid that causes insulin resistance (Fig. 7B).

**Figure 7.**
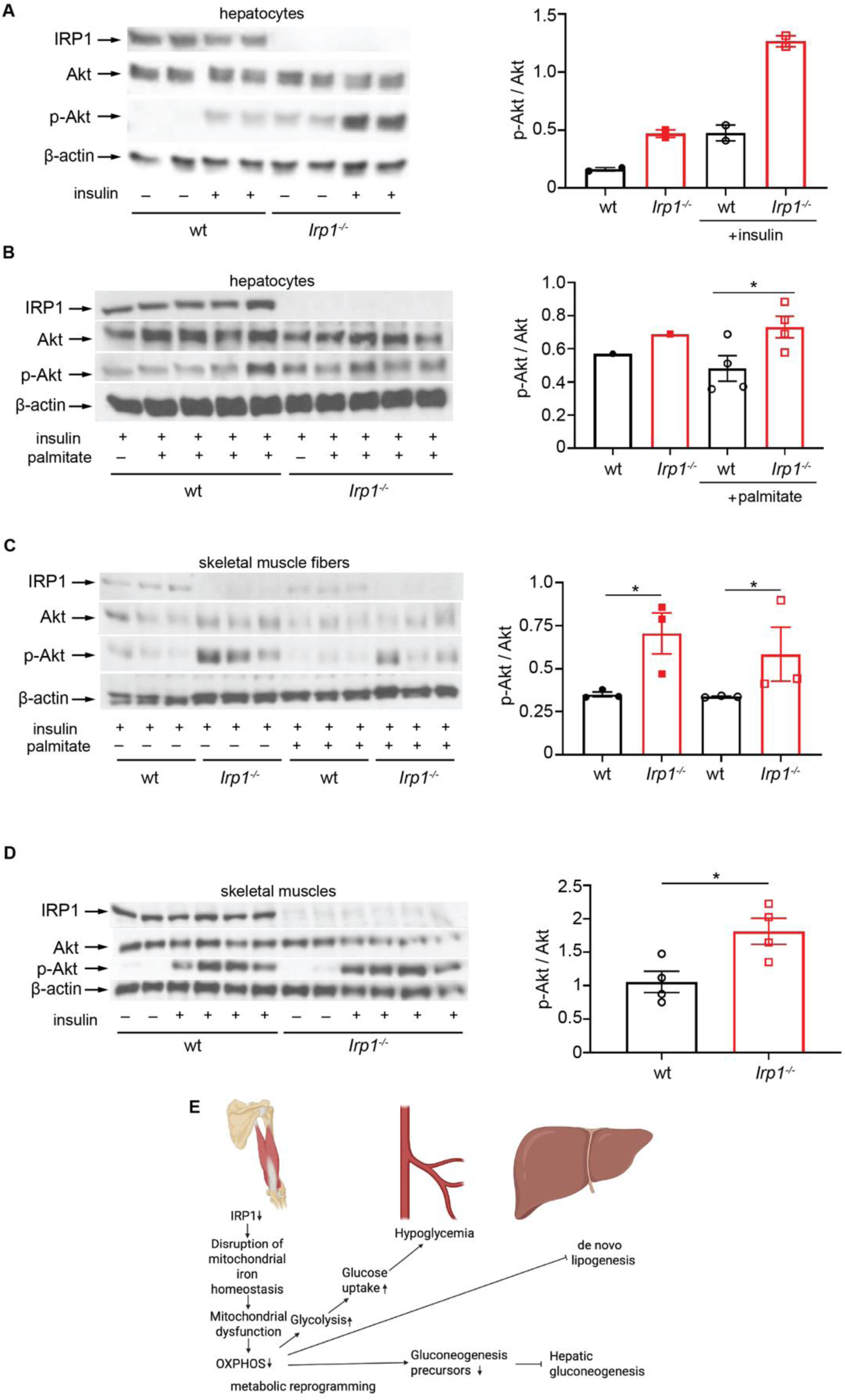
IRP1 deficiency stimulates insulin signaling. (A) Western blot analysis of phospho-Akt, Akt, IRP1 and β-actin in primary hepatocytes from *Irp1^-/-^* and wild type mice either left untreated, or previously treated with 10 nM insulin for 5 min. (B) Western blot analysis of phospho-Akt, Akt, IRP1 and β-actin in primary hepatocytes from *Irp1^-/-^* and wild type mice pretreated for 24 h with 0.4 mM palmitate (conjugated to bovine serum albumin) or not, and then treated with 10 nM insulin for 5 min. (C) Western blot analysis of phospho-Akt, Akt, IRP1 and β-actin in skeletal muscle fibers from *Irp1^-/-^* and wild type mice pretreated for 24 h with 0.4 mM palmitate (conjugated to bovine serum albumin) or not, and then treated with 50 nM insulin for 20 min. (D) Male *Irp1^-/-^* and wild type mice on high-fat diet (HFD) for 10 weeks were injected with 4U/kg insulin, or not. After 5 min, the mice were sacrificed, and skeletal muscles were analyzed by Western blotting for expression of phospho-Akt, Akt, IRP1 and β-actin. (E) A model highlighting metabolic implications of IRP1 deficiency. In (A-D), all Western blots show representative samples from each condition. Phospho-Akt to Akt ratios were quantified by densitometry and plotted on the right of each Western plot. Quantitative data are presented as the mean±SEM. Statistically significant differences are indicated by * (p<0.05) or ** (p<0.01).

A similar experiment was performed with skeletal muscle fibers isolated from wild type and *Irp1^-/-^*mice (Fig. 7C). p-Akt/Akt ratios were increased in insulin-treated *Irp1^-/-^* vs wild type muscle fibers, and remained elevated after palmitate administration. To validate this finding in vivo, wild type and *Irp1^-/-^* mice on HFD were injected with insulin or not, and skeletal muscle tissue was analyzed by Western blotting. Insulin injection stimulated Akt phosphorylation in both genotypes (Fig. 7D). However, p-Akt/Akt ratios were significantly higher in *Irp1^-/-^* vs wild type skeletal muscles, again indicating increased insulin sensitivity. These data suggest that IRP1 deficiency increases insulin sensitivity and ameliorates fat-induced insulin resistance in the liver and skeletal muscles, in agreement with the metabolic phenotype of *Irp1^-/-^*mice.

## Discussion

*Irp1^-/-^* mice were initially considered to lack any discernible pathology (Meyron-Holtz *et al*, 2004). Nevertheless, further studies revealed that these animals develop polycythemia and pulmonary hypertension due to aberrant HIF2α-dependent transcriptional induction of erythropoietin in renal interstitial fibroblasts and hepatocytes, and of endothelin 1 in pulmonary endothelial cells, respectively (Anderson *et al*., 2013; Ghosh *et al*., 2013; Wilkinson & Pantopoulos, 2013a). The mechanism involves translational de-repression of the IRE-containing HIF2α mRNA due to IRP1 deficiency. HIF2α and its homologue HIF1α are sensitive to oxygen- and iron-dependent post-translational modification by prolyl-hydroxylases (PHD1, PHD2 or PHD3), which leads to their proteasomal degradation via the von Hippel-Lindau (VHL) E3 ubiquitin ligase complex (Lee *et al*, 2020). Modest HIF2α stabilization in the mouse liver (by acute deletion of *Phd3*) has been associated with improved insulin sensitivity and relative protection against type 2 diabetes following induction of *Irs2*, another HIF2α target gene (Taniguchi *et al*., 2013; Wei *et al*., 2013).

Consistent with these data, we observed that *Irp1^-/-^*mice exhibit more pronounced insulin responsiveness and glucose clearance compared to wild type littermates, even after HFD feeding (Fig. 1). These responses were likewise associated with a trend for *Irs2* induction in the liver, arguing for potential HIF2α involvement. A modest HIF2α induction due to IRP1 deficiency could also contribute to suppressed *Srebp1*-induced lipogenesis and protection of *Irp1^-/-^* mice against liver steatosis (Taniguchi *et al*., 2013) (Fig. 2). Another argument supporting a role of HIF2α is provided by the correction of fasting hypoglycemia in older *Irp1^-/-^*mice, similar to polycythemia (Wilkinson & Pantopoulos, 2013a), and by the abrogation of this effect under HIF2α-stabilizing (Schwartz *et al*, 2019) dietary iron restriction (Figs. 1D and 5G). However, expression of gluconeogenic genes was not suppressed in *Irp1^-/-^* mice (Figs. 1I-J and S1G), despite increased insulin sensitivity. This finding also contrasts data in mouse models of modest or more severe hepatocyte-specific HIF2α stabilization due to acute ablation of *Phd3* (Taniguchi *et al*., 2013; Wei *et al*., 2013) or constitutive ablation of *Vhl* (Ramakrishnan *et al*, 2016), respectively. Both approaches led to inhibition of gluconeogenic gene expression via either stimulation of insulin signaling (Taniguchi *et al*., 2013; Wei *et al*., 2013), or repression of glucagon signaling (Ramakrishnan *et al*., 2016). Moreover, constitutive HIF2α upregulation in *Vhl*-deficient hepatocytes is known to cause severe steatosis in adult mice on a standard chow (Rankin *et al*, 2009), while *Irp1^-/-^* mice were protected against liver steatosis even following HFD feeding (Fig. 2).

Our initial data suggested a more complex phenotype for *Irp1^-/-^*mice that cannot be fully explained by molecular responses of modest HIF2α upregulation in hepatocytes, and may also be linked to energy deprivation and involve other tissues. In fact, hypoglycemia of *Irp1^-/-^* mice on standard diet was severe and only comparable to models of hepatic HIF2α overexpression above a metabolic favorable threshold (Taniguchi *et al*., 2013). We speculate that this accounts for the 18% lower survival that we noted for these animals at the age of 4 weeks. Embryonic lethality has been reported for mice with a βgeo gene trap construct inserted into the *Irp1* locus, but not for *Irp1^-/-^* mice (Galy *et al*, 2005a). The differences may be related to the genetic background of the animals, dietary factors, and possibly also the ∼4.5 times lower number of mice genotyped in the previous study (Galy *et al*., 2005a).

The liver proteomics analysis (Fig. 3) provided further hints for altered energy metabolism in *Irp1^-/-^*mice. It uncovered potential mitochondrial dysfunction and changes in expression of proteins involved in oxidative phosphorylation and energy metabolism. We corroborated these data by Western blot analysis of respiratory chain proteins (Fig. 3E) and by assessment of mitochondrial respiration in MEFs, primary hepatocytes and differentiated myotubes from *Irp1^-/-^* mice (Figs. 4 and S7). All these cell types manifested defective mitochondrial respiration and a switch to aerobic glycolysis for energy metabolism, also known as “Warburg effect” in cancer cells (Vander Heiden *et al*, 2009), which was accompanied by increased proliferation rates of MEFs and myoblasts (Fig. S8B-C). The phenotype was more pronounced in MEFs and differentiated myotubes. Skeletal muscle fibers from *Irp1^-/-^* mice exhibited increased glucose uptake, consistent with the high need for nutrients for energy production via less efficient aerobic glycolysis (Vander Heiden *et al*., 2009). Importantly, differentiated myotubes displayed aberrant oxygen consumption with palmitate as substrate, indicating defective β-oxidation (Fig. 4F), which correlated with relative protection against insulin resistance (Fig. 7C). These data support the conclusion from an earlier study that “dietary fat is less damaging to skeletal muscle metabolic function under conditions of constrained β-oxidation” (Koves *et al*, 2008).

We provide evidence that mitochondrial dysfunction is caused by disruption of iron redox balance in this organelle due to lack of IRP1 (Fig. 5). Our findings offer an additional link between iron and intermediary metabolism (Fillebeen *et al*, 2020). HFD feeding has been associated with responses to iron starvation in the liver, such as upregulation of TfR1 expression (Padda *et al*., 2015), possibly via IRP1 (Dongiovanni *et al*., 2015). These data are largely validated in Fig. 5A. Even though we did not directly compare changes in mitochondrial iron content in response to HFD, we speculate that increased supply of metabolically active iron to mitochondria is needed to cope with the metabolic stress imposed by long-term exposure to HFD. Another limitation of our study is that we did not validate the mitochondrial dysfunction data from primary cell cultures using isolated tissue mitochondria. Conversely, we did not validate the iron speciation data from tissues using primary cell cultures.

The critical role of iron is also underlined by the observations that dietary iron restriction maintained hypoglycemia (Fig. 1D), efficient glucose clearance (Fig. 5G) and poor gluconeogenesis (Fig. 5H) in *Irp1^-/-^* mice, and the glucose phenotype was partially reversed by iron dextran injection (Fig. 5I). These findings are in tune with previous studies showing that iron deficiency causes mitochondrial dysfunction and uncoupling of oxidative phosphorylation in the liver (Masini *et al*, 1994), impairs the gluconeogenic capacity of isolated hepatocytes (Klempa *et al*, 1989), enhances glucose catabolism in skeletal muscles (Henderson *et al*, 1986) and increases peripheral insulin responsiveness (Borel *et al*, 1993; Farrell *et al*, 1988). Moreover, they are consistent with the protective effects of iron deficiency and mitochondrial dysfunction against insulin resistance caused by a HFD in rats (Han *et al*, 2011).

Our data suggest that not only iron content, but also iron redox balance is critical for mitochondrial function. Notably, the loss of IRP1, a cytosolic protein, disrupted iron redox balance in mitochondria but not in the cytosol (Figs 5D and 5F). In liver mitochondria, *Irp1^-/-^* mice exhibited an increase in the Fe²⁺/Fe³⁺ ratio, primarily driven by elevated Fe²⁺ concentrations. Excess Fe²⁺ may represent a metabolically inert iron pool that fails to enter bioenergetic pathways, potentially increasing redox vulnerability. Contrary, in skeletal muscle mitochondria from *Irp1^-/-^* mice, the Fe²⁺/Fe³⁺ ratio was lower due to decreased Fe²⁺ and elevated Fe³⁺ levels, pointing to oxidative sequestration and reduced availability of metabolically active Fe²⁺. In both cases redox imbalance of iron causes mitochondrial dysfunction.

Our data highlight the important role of IRP1 in mitochondrial iron metabolism, especially under stress, and corroborate previous findings in diverse settings. For instance, *Fxn^Alb^* mice with hepatocyte-specific frataxin disruption exhibited mitochondrial dysfunction that was exacerbated upon crossing with *Irp1^-/-^* mice due to defective mitochondrial iron import (Martelli *et al*, 2015). In another example, targeted hepatocyte-specific ablation of both IRP1 and IRP2 caused liver failure and death in mice due to mitochondrial iron deficiency and dysfunction (Galy *et al*., 2010); these data also imply a contribution of IRP2. Indeed, MEFs from either *Irp1^-/-^* or *Irp2^-/-^* mice had compromised mitochondrial Fe-S cluster biogenesis due to repressed frataxin and IscU expression (Li *et al*., 2018), while IRP2 deficiency triggered a switch from oxidative phosphorylation to aerobic glycolysis in MEFs (Li *et al*, 2019). However, contrary to the metabolic phenotype of *Irp1^-/-^* mice reported herein, *Irp2^-/-^* animals rather exhibit glucose intolerance and develop diabetes (Santos *et al*, 2020). Thus, the systemic metabolic functions of IRP1 and IRP2 are distinct. Dissecting the mechanism by which IRP1 modulates mitochondrial iron metabolism awaits further investigation.

Our metabolomics analysis uncovered extensive metabolic reprogramming in the liver and skeletal muscles of *Irp1^-/-^* mice, presumably due to mitochondrial dysfunction and inefficient energy production. These responses were exacerbated under conditions of high energetic needs following metabolic stress (HFD) or dietary iron restriction (Figs. 6 and S10-11), and are reminiscent of metabolic changes reported in *Tfr1^mu/mu^*mice, bearing skeletal muscle-specific disruption of TfR1 (Barrientos *et al*, 2015). In this animal model, TfR1 deficiency in skeletal muscles caused severe iron deficiency incapacitating mitochondrial energy production that led to growth arrest and a muted attempt to switch to fatty acid β-oxidation, consuming fat stores. Even though hepatic gluconeogenesis was stimulated, amino acid substrates became limiting and lethal hypoglycemia developed shortly after birth.

In *Irp1^-/-^* mice on IDD or HFD, amino acid substrates likewise became limiting for gluconeogenesis, which might also have been impaired by energy deprivation. Paradoxically, insulin signaling was stimulated in these animals rather than suppressed (Fig. 7). This very likely contributed to IDD-induced hypoglycemia or protection against HFD-induced hyperglycemia, and increased glucose disposal by skeletal muscles. The protection against HFD-induced hyperglycemia was more prominent in male mice (Figs. 1E and S1F). The molecular basis for the apparent sexual dimorphism is not clear and will be addressed in follow-up studies.

On a final note, *Irp1^-/-^* mice were initially reported to lack any discernible pathology (Meyron-Holtz *et al*., 2004). Nevertheless, we and others demonstrated that these animals develop polycythemia due to translational de-repression of renal HIF2α mRNA and subsequent transcriptional induction of its downstream target erythropoietin (Anderson *et al*., 2013; Ghosh *et al*., 2013; Wilkinson & Pantopoulos, 2013a). The physiological relevance of our findings was subsequently validated by a meta-analysis of genome-wide association studies (GWAS) involving 684,122 individuals from Iceland and UK, which identified the IRP1-encoding *ACO1* gene as a major homeostatic regulator of hemoglobin concentration (Oskarsson *et al*, 2020). We expect that the metabolic phenotype of *Irp1^-/-^* mice reported herein will spark analogous validation studies.

In conclusion, our data uncover a novel function of IRP1 as a metabolic switch (Fig. 7E), and provide evidence that this involves its iron regulatory activity. Moreover, they suggest that targeting IRP1 may offer therapeutic benefits against hyperglycemia, insulin resistance and metabolic dysfunction associated steatotic liver disease (MASLD), the hepatic manifestation of the metabolic syndrome (Targher *et al*, 2024).

## Methods

### Animals

*Irp1^-/-^* mice on C57BL/6 background (Galy *et al*., 2005a) were kindly provided by Dr. M. W. Hentze (EMBL, Heidelberg, Germany). *Irp1^+/-^* mice were generated by breeding *Irp1^-/-^* with congenic *Irp1^+/+^* (wild type) mice. Breeding pairs from these animals were used to generate *Irp1^+/+^*(wild type) and *Irp1^-/-^* littermates that were used throughout this study. The animals were housed in macrolone cages (up to 5 mice per cage, 12:12-hour light-dark cycle: 7:00AM to 7:00PM; 22±1°C, 60±5% humidity) according to institutional guidelines. Where indicated, the mice were fed a high-fat diet (HFD with 43% fat; ssniff Spezialdiäten GmbH S9552-E034) or its equivalent control diet (with 12% fat; ssniff Spezialdiäten GmbH S9552-E033). In other experiments, the mice were fed an iron-deficient diet (IDD; TD.80396 containing 2-6 ppm iron) or its equivalent control diet (Teklad Global 18% protein 2918 containing 200 ppm iron). All experimental procedures were approved by the Animal Care Committee of McGill University (protocol 4966).

### Cell culture

Mouse embryonic fibroblasts (MEFs) were obtained from wild type and *Irp1^-/-^* mouse embryos. Cells were cultured in Dulbecco’s modified Eagle’s medium (DMEM) supplemented with 10% heat-inactivated fetal bovine serum (Wisent), non-essential amino acids, 100 U/ml penicillin and 100 μg/ml streptomycin, and immortalized using a pBABE-neo large T antigen cDNA (Addgene #1780). Primary hepatocytes were prepared from adult mice using a two-step collagenase perfusion technique and cultured as described (Fillebeen *et al*, 2018). Primary myoblasts were prepared from 4 to 6-week-old mice as described (Lazure *et al*, 2023). Differentiation to myotubes was in DMEM supplemented with 5% horse serum for 5 days. Muscle fibers were isolated from soleus muscle (Sahinyan *et al*, 2022) and cultured in DMEM supplemented with 10% fetal bovine serum and antibiotics.

### Glucose, pyruvate, and insulin tolerance tests

*Irp1^-/-^* mice and wild type littermates were fasted for 4-16 hours before the experiments. The animals were intraperitoneally injected with 1 g/kg glucose for glucose tolerance test (GTT), 2 g/kg sodium pyruvate for pyruvate tolerance test (PTT), or 0.5 U/kg insulin (Humulin, DIN 00586714) for insulin tolerance test (ITT). Where indicated, mice were intraperitoneally injected with iron dextran (Sigma) at 15 mg/kg, three times with 2 h intervals; the last injection was performed 24 h before the GTT. Blood glucose levels were measured at t=0, 15, 30, 60, and 120 min using the OneTouch Verio Flex® blood glucose meter.

### Glucose uptake assay

Isolated muscle fibers were starved in glucose deficient DMEM for 45 minutes before the assay. Subsequently, the media were replaced with DMEM containing 4.5 g/L glucose and supplemented with 1% fetal bovine serum and antibiotics, and the muscle fibers were incubated for 6 h. Culture supernatants (100 μl) were collected at t=0, 3 and 6 h, and glucose was measured by using glucose oxidase liquid reagents (Pointe Scientific G7521120). Glucose uptake was calculated as the difference of glucose amount collected at t=0 and the specific time points. Data was normalized to total protein.

### Seahorse experiments

Mitochondrial respiration and glycolysis were analyzed in MEFs, primary hepatocytes, or differentiated myotubes by the Seahorse assay (Divakaruni & Jastroch, 2022). Details are provided in the Supplemental Methods section.

### Lactate assay

Cells were cultured in DMEM supplemented with 1% fetal bovine serum and antibiotics (100 U/ml penicillin, 100 μg/ml streptomycin). Old media were replaced with fresh, and aliquots (100 μl) were collected at t=3, 6 and 24 h. Lactate was measured using a lactate oxidase reagent kit (Pointe Scientific L7596-50). Data were normalized to total protein.

### Isolation of cytosolic and mitochondrial fractions

Liver and skeletal muscles from *Irp1^-/-^* mice and wild type littermates were processed to separate mitochondrial and cytosolic fractions by using a mitochondria isolation kit (Thermo Fisher #89801).

### Iron redox speciation analysis

For redox speciation analysis of Fe^2+^ and Fe^3+^, the method outlined in (Solovyev *et al*, 2017) was significantly modified to reduce the overall analysis time (including analysis and column cleaning) and to optimize detection for ICP-KED-MS (Inductively Coupled Plasma - Kinetic Energy Discrimination - Mass Spectrometry). Details are provided in the Supplemental Methods section.

### Quantitative real-time reverse transcription PCR (qPCR)

Total RNA was extracted from tissues by using the RNeasy kit (Qiagen). Purity was assessed by 260/280 nm absorbance ratios and quality was monitored by agarose gel electrophoresis. cDNA was synthesized from 1 μg RNA by using the OneScript Plus cDNA Synthesis Kit (Applied Biological Materials). qPCR was performed in a 7500 Fast Real Time PCR System (Applied Biosystems) with gene-specific primers provided in the “Reagent and resource” file. Primer pairs were validated by dissociation curve analysis and demonstrated amplification efficiency between 90% and 110%. SYBR Green (Bioline) and primers were used to amplify products under following cycling conditions: initial denaturation 95°C 10 min, 40 cycles of 95°C 5 s, 58°C 30 s, 72°C 10 s, and final cycle melt analysis between 58°-95°C. Relative mRNA expression was calculated by the 2^-ΔΔCt^ method (Livak & Schmittgen, 2001). Data were normalized to murine ribosomal protein L19 (*Rpl19*).

### Western blotting

Western blot analysis was performed as earlier described (Katsarou *et al*, 2021). Antibodies and dilutions are provided in the “Reagent and resource” file. Immunoreactive bands were quantified by densitometry with the ImageJ software.

### Histology

Liver specimens from 3 mice per experimental group were fixed in 10% buffered formalin and embedded in paraffin. Samples were cut at 4-µm, placed on SuperFrost/Plus slides (Fisher) and dried overnight at 37°C. De-paraffinized slides were used for H&E or Periodic Acid-Schiff (PAS) staining.

### Statistical analysis

Quantitative data were expressed as mean ± standard error mean (SEM). Statistical analysis was performed by using the GraphPad Prism 10.5.0. Comparisons between two independent groups were done by the unpaired Student’s t-test. A p value of less than 0.05 was considered statistically significant.

### Lipidomics, proteomics, metabolomics studies and other assays

Experimental outline and details are provided in the Supplemental Methods section.

## Acknowledgments

We thank Dr. M. W. Hentze (EMBL, Heidelberg, Germany), Dr. Bruno Galy (DKFZ, Heidelberg, Germany) and the late Dr. G. D. Lopaschuk (University of Alberta, Edmonton, Canada) for stimulating discussions in the early phase of this work. We also thank Dr. Naciba Benlimame for assistance with histology, Veronique Michaud and Xiaohong Liu for support with animal procedures, and Dr. Maia Kokoeva for providing access to metabolic cages. This work was supported by a grant from the Canadian Institutes of Health Research (CIHR; PJT-186193). The work of VV was financially supported by the Deutsche Forschungsgemeinschaft (DFG) through the Priority Program “Ferroptosis: from Molecular Basics to Clinical Applications” (SPP 2306).

## Author Contributions

<u>Wen Gu</u>: Investigation, formal analysis, methodology, writing original draft. <u>Nicole Wilkinson</u>: Investigation. <u>Carine Fillebeen</u>: Formal analysis, methodology, project administration. <u>Darren Blackburn</u>: Methodology. <u>Korin Sahinyan</u>: Methodology. <u>Eric Bonneil</u>: Methodology, formal analysis. <u>Tao Zhao</u>: Investigation. <u>Zhi Luo</u>: Formal analysis. <u>Vahab Soleimani</u>: Methodology. <u>Vincent Richard</u>: Methodology, formal analysis. <u>Christoph H. Borchers</u>: Methodology. <u>Albert Koulman</u>: Methodology. <u>Benjamin Jenkins</u>: Methodology. <u>Bernhard Michalke</u>: Methodology. <u>Hans Zischka</u>: Methodology. <u>Judith Sailer</u>: Methodology. <u>Vivek Venkataramani</u>: Methodology. <u>Othon Iliopoulos</u>: Methodology, conceptualization. <u>Gary Sweeney</u>: Methodology, resources, conceptualization. <u>Kostas Pantopoulos</u>: Conceptualization, supervision, funding acquisition, writing review and editing.

## The Paper Explained

Iron regulatory protein 1 (IRP1) is known for its role in cellular iron metabolism, but its impact on whole-body energy regulation was previously unclear. This study reveals that loss of IRP1 in mice leads to mitochondrial dysfunction, shifts cellular energy production from oxidative phosphorylation to glycolysis, and enhances insulin sensitivity. As a result, IRP1-deficient mice are protected from high-fat diet-induced hyperglycemia and liver steatosis—hallmarks of metabolic syndrome. These findings identify IRP1 as a key metabolic regulator that links iron homeostasis to mitochondrial function and glucose metabolism. Targeting IRP1 or its downstream pathways may offer new therapeutic avenues for managing insulin resistance and metabolic liver disease.

